# Are increasing honey bee colony losses attributed to *Varroa destructor* in New Zealand driven by miticide resistance?

**DOI:** 10.1101/2023.03.22.533871

**Authors:** Rose A. McGruddy, Mariana Bulgarella, Antoine Felden, James W. Baty, John Haywood, Philip Stahlmann-Brown, Philip J. Lester

## Abstract

The most devastating pest to honey bees (*Apis mellifera*) worldwide is the parasitic mite *Varroa destructor*. The development of miticide-resistant mite populations has been a major driver of colony loss in many countries. We investigated the threat Varroa poses to honey bee populations in New Zealand and tested the effectiveness of the two most popular chemical treatments used by beekeepers. Colony losses reported by New Zealand beekeepers have risen over five consecutive years from 2017 to 2021, as have the proportion of losses attributed to Varroa, with this parasite found to be the main driver of colony loss in 2021. Varroa resistance to miticide treatments flumethrin and amitraz was tested. The concentration of flumethrin required to kill 50% of the mites (LC_50_) was 156 μg/g, 13 times greater than the adjusted LC_50_ value of 12 μg/g observed in a trial also conducted in New Zealand in 2003, thus indicating evidence of developing mite resistance to flumethrin in New Zealand. Molecular analyses searching for mutations in the Varroa genome known to be associated with flumethrin resistance found no evidence of such mutations, suggesting that any extant resistance to flumethrin has evolved independently in New Zealand. No evidence of resistance to amitraz was found, as the LC_50_ value of 12 μg/g was lower than what was observed in the 2003 trial (110 μg/g). Further development of integrated pest management, such as gene-silencing RNA interference (RNAi) and selective breeding of Varroa-resistant bees, is needed to effectively manage a parasite that threatens global agriculture.

## Introduction

The western honey bee, *Apis mellifera*, plays a critical role in crop pollination across the globe (Aizen and Harder 2009). Reliance on pollinator-dependent crops has risen over recent decades (Aizen *et al*. 2019) and there is evidence of pollinator limitation in many fruit and vegetable crops as honey bee stocks struggle to keep up with demand (Breeze *et al*. 2014, Reilly *et al*. 2020). The loss of pollination services not only threatens food security, but also poses a significant risk to economies that depend on the international trade of pollinated crops (Eilers *et al*. 2011, Vanbergen and Insect Pollinators Initiative 2013, Murphy *et al*. 2022). The deficit between the need for pollinators and the availability of bee colonies is due in part to a decline in honey bee health and increases in hive mortality rates (Dolezal *et al*. 2019, Steinhauer *et al*. 2021). Stressors such as parasites, pesticides and disease are recognised drivers of colony loss (Goulson *et al*. 2015, Di Prisco *et al*. 2016). The most significant threat to the apiculture industry worldwide is frequently cited as the ectoparasite *Varroa destructor* (Rosenkranz *et al*. 2010, Traynor *et al*. 2020, Jack and Ellis 2021). This parasite feeds on the fat body of bees, a tissue crucial to bee health that is involved in immune function, the detoxification of pesticides and winter survival (Amdam *et al*. 2004, Ramsey *et al*. 2019). Varroa is also a vector for honey bee viruses, particularly the Deformed wing virus (DWV) (Wilfert *et al*. 2016), which is associated with a wide range of clinical symptoms including crippled wings, cognitive impairment and reduced lifespan (Dainat *et al*. 2012, Francis *et al*. 2013). Both Varroa and DWV are strong predictors of colony collapse over winter (Dainat *et al*. 2012, Francis *et al*. 2013). Colonies that are not treated for Varroa typically collapse within three years (Rosenkranz *et al*. 2010).

The most widely used Varroa control methods currently on the market are synthetic or organic miticides. These chemicals are usually applied as impregnated strips which are placed within the beehive and left for a period of four to ten weeks. Of the synthetic miticides, pyrethroid and formamidine based treatments have historically been popular choices for mite control. Pyrethroids specifically act on voltage-gated sodium channels, prolonging the opening of channels, causing paralysis and death (Narahashi 2000, Wang *et al*. 2003). Formamidines are designed to target and stimulate alpha-2 adrenoceptors, interfering with the nervous system, which can result in a variety of outcomes, including impairment of consciousness and convulsions (Dekeyser and Downer 1994). Synthetic miticides are selected and developed based on their ability to control Varroa without killing their host.

A key issue with miticides is the development of resistance by Varroa to many products on the market (Hernández-Rodríguez *et al*. 2021). Resistance to pyrethroid-based treatments was first reported from Italy around 1991 (Martin 2004), and since then, pyrethroid-resistant mites have been found in the United Kingdom, Europe, Canada and USA (Martin 2004, Mitton *et al*. 2022). Mutations have been associated with mite resistance to pyrethroids in Varroa and other arthropods (González-Cabrera *et al*. 2013, González-Cabrera *et al*. 2016, Millán-Leiva *et al*. 2021). In Varroa, amino acid substitutions in the voltage-gated sodium channel protein at position 925 in particular have been observed in a number of mite populations that demonstrated resistance (Millán-Leiva *et al*. 2021). Incidences of resistance to formamidine-based Varroa treatments have been less common, although inefficacy has been detected in some studies (Thompson *et al*. 2002, Kamler *et al*. 2016, Higes *et al*. 2020, Rinkevich 2020, Hernández-Rodríguez *et al*. 2021). A growing number of treatments appear to be becoming ineffective against Varroa mites, with a lack of effective alternatives available to replace them.

Varroa was first discovered in New Zealand in the year 2000 (Goodwin 2004). Pyrethroids were the first treatments to be registered for use against Varroa in 2000 and 2001, followed by formamidine treatments in 2004. As resistance had already been reported in numerous countries by this time, a study examining miticide efficacy on Varroa populations in New Zealand was conducted in 2003 by Goodwin *et al*. (2005). Their study did not find sufficient evidence to conclude that mites were resistant to either flumethrin (a pyrethroid-based treatment) or amitraz (formamidine-based treatment) and beekeepers have continued to use these treatments to control Varroa.

One management tool that has proven useful in identifying the threats to honey bees has been the establishment of a survey whereby beekeepers can report colony losses each winter. Colony loss surveys were first implemented in Canada in 2003 following reports of Varroa resistance to treatments (Currie *et al*. 2010). In the years following, other regions of the world including the USA, Canada, Europe, Asia, the Middle East and Africa began conducting surveys in the wake of increased levels of overwintering colony losses (vanEngelsdorp *et al*. 2008, van der Zee *et al*. 2012, Brodschneider *et al*. 2018). In 2008, a standardised survey, now known as the “COLOSS” survey, was developed to make colony loss data comparable internationally (Gray *et al*. 2022). These surveys enable spatial and temporal analyses of the threats to honey bees. A survey based on the COLOSS questionnaire was first conducted in New Zealand in 2015 and has been undertaken annually since (Brown *et al*. 2018). In addition to overall losses, the New Zealand survey has measured beekeepers’ attributions of losses, including to parasitic Varroa mites and related complications, since 2017. The annual survey also includes detailed questions on Varroa monitoring and treatment.

The aims of this study were to 1) report the role that Varroa has played in colony losses in New Zealand according to beekeepers that responded to the survey, 2) describe current Varroa management strategies practiced by commercial beekeepers and any changes in this strategy over a five-year period and 3) test for evidence of Varroa resistance to the two most commonly utilised chemical treatments in New Zealand: flumethrin and amitraz.

## Materials and Methods

### Varroa and colony losses in New Zealand

The New Zealand Colony Loss Survey covers topics related to hive management, such as the number of colonies lost over winter, beekeepers’ attributions of losses (including Varroa and related complications) and Varroa control methods used. The current study analysed survey results for the years 2017-2021 only, as “*suspected Varroa and related complications*” were not included as an attributable cause of losses prior to 2017. In the 2021 survey, beekeepers were also given the opportunity to provide feedback on the perceived effectiveness of the treatments they used to combat Varroa, to give insight into the effectiveness of current treatments and to detect any early signs of developing resistance.

New Zealand beekeepers are legally obligated to register their hives under the Biosecurity Act 1993 (MPI 1993), and all registered beekeepers were invited to participate in the online colony loss survey. This mandatory registration also allows for the percentage of beekeepers that participated in the survey to be estimated. In 2017, 2,066 beekeepers completed the survey, a response rate of 30.9% of all beekeepers nationwide. In the years following, the number of beekeepers that participated were 3,655 (42.3%), 3,456 (36.7%), 2,863 (32.0%) and 4,355 (49.1%) for the years 2018, 2019, 2020 and 2021, respectively (Stahlmann-Brown *et al*. 2021). Our investigation into Varroa management strategies differed from previous work by only focusing on responses from commercial beekeepers only, defined here as having more than 350 hives at the beginning of winter (according to the definition by New Zealand’s Ministry for Primary Industries (MPI 2020)). These beekeepers manage the majority of hives in the country and therefore their success in controlling Varroa is of the greatest economic interest. Loss rates from the survey were calculated using standard approaches for estimating colony losses (van der Zee *et al*. 2013) as detailed in Stahlmann-Brown and Robertson (2022). Statistical analyses were conducted in R 4.2.0 (R Development Core Team 2020). Overall loss rates and corresponding confidence intervals (CI) were calculated with a quasi-binomial generalised linear model and logit link function. A test of equal or given proportions (“prop.test”) determined if there was a significant change in the overall loss rates or losses attributed to Varroa over the five-year period of 2017-2021.

### Testing for pesticide resistance in mites

Trials were conducted in April and May of 2022 at Victoria University of Wellington, Wellington, New Zealand. The testing protocol from Goodwin *et al*. (2005) was followed to allow for comparison. Analytical standard grade flumethrin (Sigma Aldrich/Merck, New Zealand) and amitraz (AK Scientific, USA), were diluted in hexane (Sigma Aldrich/Merck, New Zealand). Flumethrin was tested at 0, 10, 20, 40, 80, 160, 320 and 640 μg/g. Amitraz was tested at 0, 2, 5, 10, 25, 50, 100, 200 and 400 μg/g. Petri dishes were prepared with the different concentrations following Goodwin *et al*. (2005) methodology with some minor modifications, as follows. Fully-refined paraffin wax (58°C melting point, National Candles Ltd., New Zealand) was melted in a microwave and 50 mL was poured into wide-mouth, graduated bottles and weighed. Twenty-five mL of the respective pesticide concentration in hexane was added to the bottles and left in a hot water bath at 60°C for approximately 8 hours until the hexane evaporated, and each bottle returned to the original weight. The mixture of paraffin and pesticide was then swirled and poured into four 35-mm sterile petri dishes (Corning, USA) to a depth of ~4 mm and kept in a fridge until use.

Mites were collected from hives at Victoria University of Wellington that had not been treated for Varroa for six months. The freshly collected mites were counted out into groups of approximately 20. Each group of mites was transferred to petri-dishes containing the treatment (either flumethrin or amitraz of a particular concentration, or a control treatment of plain paraffin), where they were left for one hour. The mites were then transferred to a third dish of the same size, along with 2-3 bee pupae collected from the same colony as the mites and placed in an incubator at 32-34°C, with 50% relative humidity (RH) for 48 hours until the survival assessment (Figure S1, Supplemental material). Analysis of the data was conducted using SPSS 28 (Akçay 2013). Abbott’s correction was used to account for mite mortality in the controls for both the Goodwin *et al*. (2005) and the 2022 datasets. The adjusted proportion of dead mites for each study was then fitted using a probit regression model on concentration (log scale). The adjusted Lethal Concentration at 50% (LC50) and associated 95% CIs were then estimated for the flumethrin and amitraz treatments.

### Investigating mutations associated with pesticide resistance in mites

We investigated two specific amino acid residue substitutions located on the *Varroa destructor* pyrethroid susceptible sodium channel (Na) gene (GenBank accession number KC152655), at nucleotide positions 1689-1691 (residue substitution M918L) (Rinkevich *et al*. 2013) and 1710-1712 (residue substitution L925V/M/I) known to be associated with flumethrin resistance in Varroa (González-Cabrera *et al*. 2013, González-Cabrera *et al*. 2016, Millán-Leiva *et al*. 2021). We aligned Varroa RNA-Seq reads obtained in another study (Lester *et al*. 2022) onto the KC152655 FASTA file using HISAT 2.0 with default parameters (Kim *et al*. 2015). The resulting BAM files were visually inspected in Geneious 11.1.5 (Kearse *et al*. 2012) to check for nucleotide polymorphisms.

In order to further investigate the presence of the *Varroa destructor* pyrethroid susceptible sodium channel (Na) gene (GenBank accession number KC152655) for mutations at positions 1710-1712, 10 mite samples were taken from locations throughout the country, including mites from the experimental hives in Wellington (Supp. Table 1, Supplemental material). Each individual mite was placed in a 2 mL microtube (Sarstedt, Germany). Five 3.2 mm stainless steel beads (Next Advance Inc., USA), 500 μL of GENEzol DNA Plant Reagent (Geneaid Biotech, Taiwan) and 2.5 μL of β-mercaptoethanol (Sigma Aldrich, USA) were added to the tube. Samples were homogenised for one cycle of 20 s each at 8,000 rpm in a Precellys Evolution homogeniser (Bertin, France). DNA and RNA was simultaneously isolated with a 24:1 chloroform–isoamyl alcohol mixture (BioUltra, Sigma Aldrich, USA), followed by isopropanol precipitation (Sigma Aldrich, USA), and an ethanol purification step (VWR Chemicals, UK). DNA/RNA was then eluted in 15 μL of nuclease-free water (Ambion, Life Technologies, USA), quantified using a NP80 NanoPhotometer (Implen, Germany) and kept at −80°C until use.

RNA samples (70 ng) were prepared for PCR by reverse transcription in 10 μL reactions using qScript cDNA SuperMix (Quantabio, USA). Two PCR assays were conducted on each sample. The first used primers Vd_L925V_F (5’-CCAAGTCATGGCCAACGTT-3’) and Vd_L925_R (5’-AAGATGATAATTCCCAACACAAAGG-3’), which generated 97 base pair products and were used to identify mutations at positions 1710-1712 (amino acid residue 925), developed by González-Cabrera *et al*. (2013). A second set of primers, Vd_general_407_F (5’-GGTCTGGAAGGCGTACAAGG-3’) and Vd_general_407_R (5’-TTGAGTACGACCAGGTTGCC-3’), amplified a larger product (406-407 base pairs), and were used to screen for mutations across a longer stretch of the gene. Reactions were set up with primers at 0.4 μM, 14 ng cDNA, 7.5 μL MyTaq Red (Bioline/Meridian Bioscience, USA), and water to a final volume of 15 μL. Run conditions were as follows: 95 °C for 1 min and then 35 cycles of 95 °C (15 s), 60 °C (15 s) and 72 °C (10 s). PCR products were then resolved by 2% agarose gel electrophoresis (100 V, 30 min), and visualised using SYBR Safe DNA gel stain (Invitrogen/ThermoFisher Scientific, USA). Products were then prepared for sequencing using ExoSAP-IT PCR Product Cleanup Reagent (Applied Biosystems/ThermoFisher Scientific, USA) following manufacturer guidelines. Sequencing was performed on an ABI 3130×1 Genetic Analyzer (Applied Biosystems, USA) at Massey Genome Service (Palmerston North, New Zealand). We visually inspected and aligned the forward and reverse gene sequences of the same mite using the default alignment algorithm implemented in Geneious Prime 2023.0.4 (http://www.geneious.com).

## Results

### Varroa and colony losses in New Zealand

The first goal of this study was to report losses attributed to Varroa based on the responses of beekeepers in the New Zealand Colony Loss Survey. Total colony loss rates have increased significantly over the last five years (*χ*^2^ = 3622.6, *df* = 4, *P* < 0.0001), from 9.70% [95% CI: 9.36% −10.04%] in 2017 to 13.59% [95% CI: 13.23% −14.01%] in 2021 (Figure 1). Among beekeepers who lost colonies, the proportion of losses attributed to Varroa has also increased significantly over the last five years (*χ*^2^ = 8215.5, *df* = 4, *P* < 0.0001), from 16.9% [95% CI: 15.3% −18.1%] in 2017 to 38.9% [95% CI: 37.7% −40.0%] in 2021 (Fig. 1). According to beekeepers that participated in the questionnaire survey in 2022, Varroa was the main driver of colony loss over winter 2021.

**Figure 1.**
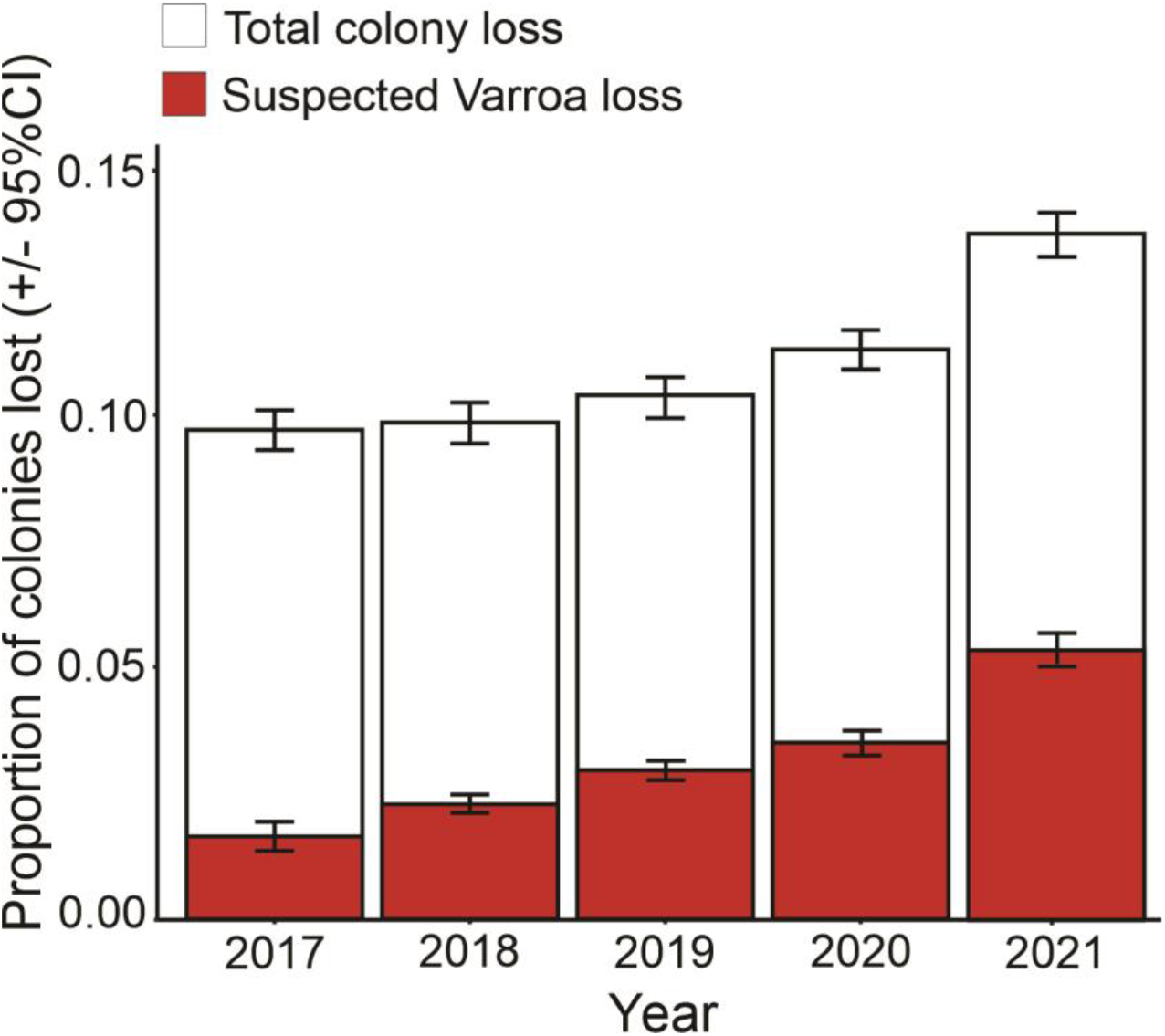
Bar graph depicting the proportion of total colonies lost (± 95% CI) and colonies that beekeepers suspected were lost due to Varroa (± 95% CI) for the years 2017-2021 in New Zealand. Results are based on reports from all beekeepers that participated in the annual New Zealand colony loss survey. The number of bee colonies reported on each year ranged between 238244-379862.

To find possible explanations for the observed increase in overall colony losses attributed to Varroa, the management strategies of commercial beekeepers were investigated. According to beekeepers that participated in the colony loss survey, amitraz and flumethrin were the two most commonly utilised Varroa treatments in New Zealand each of the five years analysed. Amitraz was the most popular choice, used annually by 85-92% of commercial beekeepers over the 2017-2021 period (Table 1). Flumethrin was used annually by 68-80% of commercial beekeepers as part of their hive treatment against Varroa over that time (Table 1). The majority of commercial beekeepers used both amitraz and flumethrin in the same year (63-75%). The other common control treatments utilised were oxalic and formic acid, which are organic miticides. The use of oxalic acid has increased steadily over the five year period, with 41.8% of beekeepers reporting its use against Varroa in 2021, whereas in 2017 only 19.5% used oxalic acid (Table 1). Formic acid use has fluctuated year to year, with 11.5-19.1% of beekeepers applying it to hives annually.

**Table 1.**
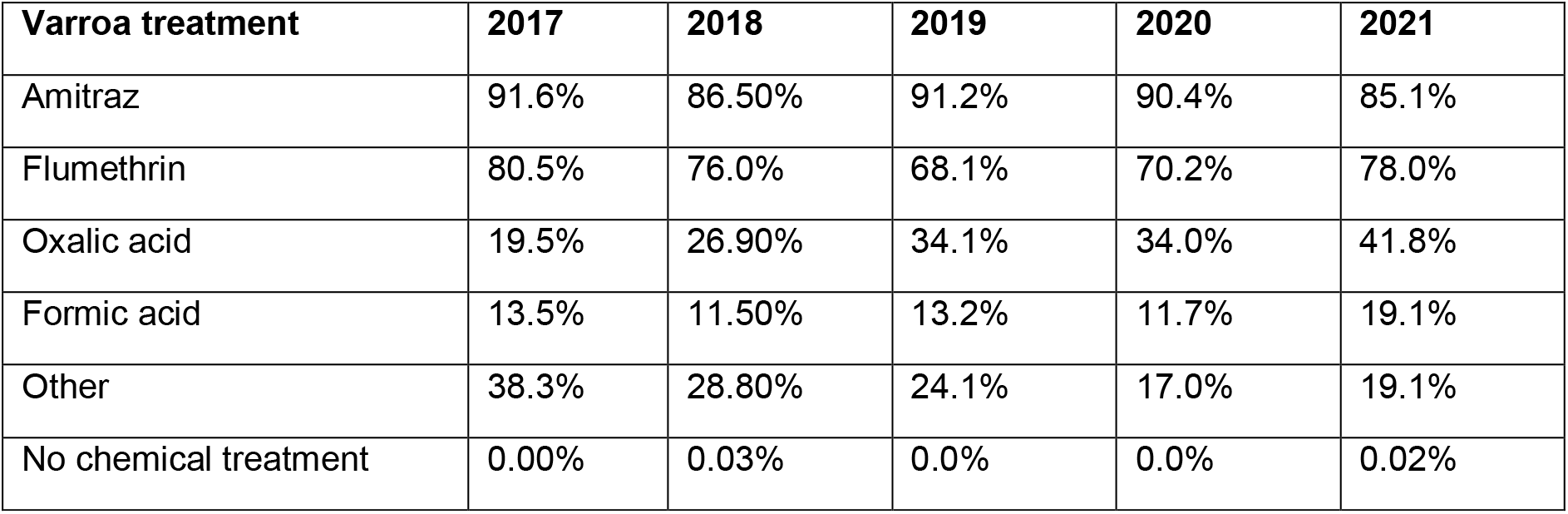
The most commonly utilised chemical treatments for Varroa as reported by commercial beekeepers in the annual colony loss survey in New Zealand from 2017-2021. The category “Other” contains control methods that aren’t already listed, such as thymol, fogging, drone brood removal and hyperthermia.

Of the beekeepers that used amitraz treatments in the 2020/2021 season, 27.6% found the treatment to be “completely successful” against Varroa, with 64.3% finding it to be “mostly successful” (Table 2). Only 8.1% reported amitraz to be either “partly” or “not at all” successful. For flumethrin, 17.9% found the treatment to be completely successful, 63.4% thought it was mostly successful and 18.8% of beekeepers reported the pyrethroid-based control to be only partly or not at all successful in controlling Varroa.

**Table 2.** Efficacy of flumethrin and amitraz, the two most commonly utilised chemical treatments for Varroa, as reported by commercial beekeepers in the annual colony loss survey in New Zealand in 2021. Efficacy in controlling Varroa was categorised as either “completely successful”, “mostly successful”, “partly successful” or “not at all successful”. The responses are displayed as a proportion of all commercial beekeepers that reported using that chemical.

| **Varroa treatment** | **Completely successful** | **Mostly successful** | **Partly successful** | **Not at all successful** |
| --- | --- | --- | --- | --- |
| Amitraz | 27.6% | 64.3% | 6.1% | 2.0% |
| Flumethrin | 17.9% | 63.4% | 16.1% | 2.7% |

### Testing for pesticide resistance in mites

The experiment testing for pesticide resistance in New Zealand populations of mites found a much higher concentration of flumethrin was required to kill mites compared to the concentration required in the 2003 study (Goodwin *et al*. 2005). The adjusted LC_50_ value for flumethrin in 2003 was 12 μg/g [95% CI = 8 - 17], whereas the adjusted LC_50_ value in 2022 had increased to 156 μg/g [95% CI = 115 - 217]. The concentration of flumethrin required to reach an average mite mortality of 50% in 2022 was a 12 fold-change higher compared to what was observed in 2003 (Figure 2). For amitraz, the adjusted LC_50_ value was 110 μg/g [95% CI = 39 – 217] in 2003 and decreased to 12 μg/g [95% CI = 10-16] in 2022. A similar concentration of amitraz was required to achieve 50% average mortality in both studies (Figure 2). We note that in the 2003 experiment there was high variability in the proportion of mites that died for the flumethrin treatment. For example, at a concentration of 1 μg/g, Goodwin *et al*. (2005) observed mortality ranging from 0-99.7% in different replicates. This level of variability was not observed in the 2022 trial. It is also worth noting that the amitraz treatment in the 2003 experiment was unable to achieve an average mortality rate above 65% at any concentration, whereas 100% mortality was achieved in the 2022 study.

**Figure 2.**
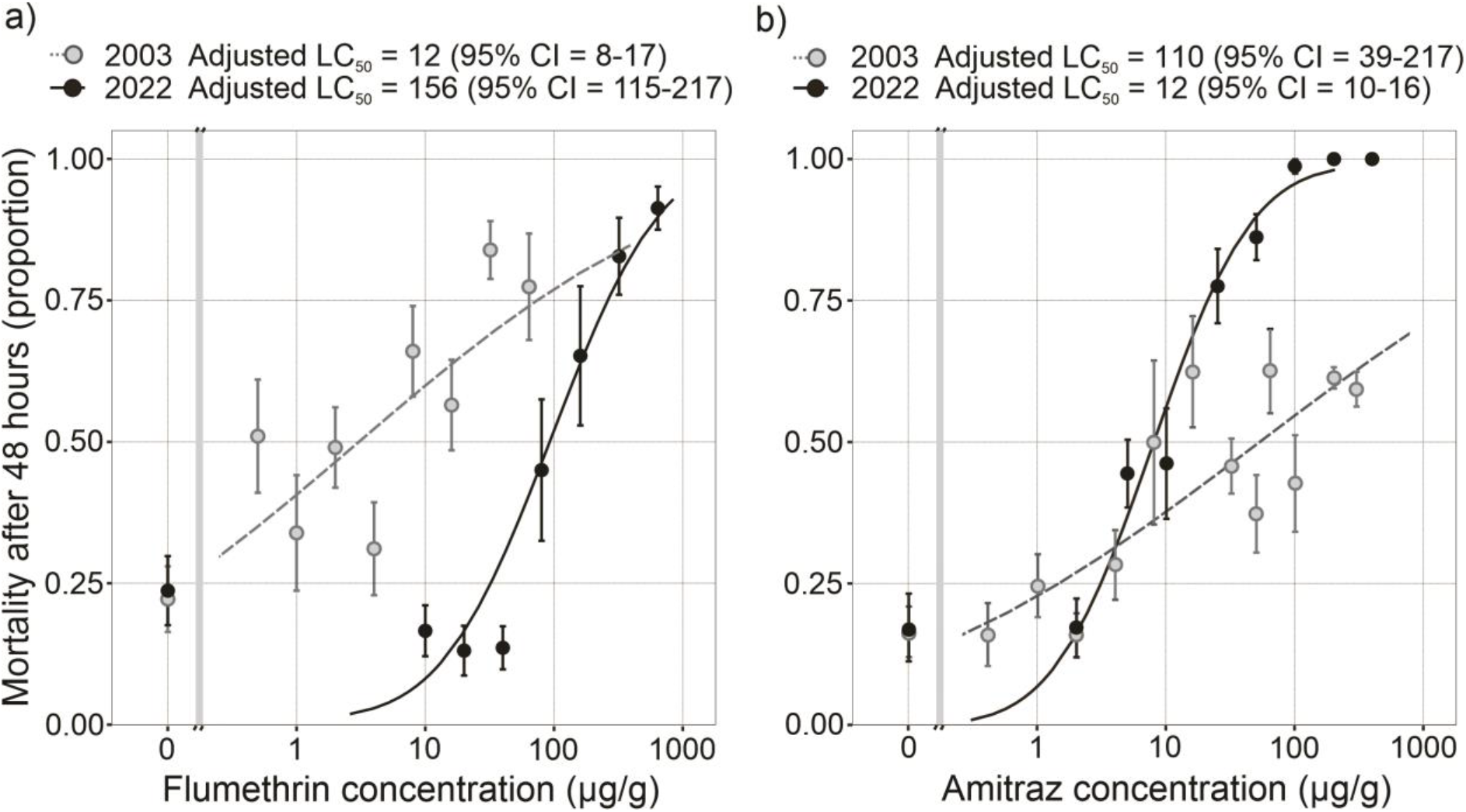
Comparison of the average proportion (±1 SE) of Varroa destructor killed at each chemical concentration in 2003 (Goodwin et al. 2005) and the current study for a) flumethrin and b) amitraz. The adjusted LC_50_ value for flumethrin in 2003 was 12 μg/g [95% CI = 8 - 17]. The adjusted LC_50_ value in 2022 was 156 μg/g [95% CI = 115 - 217]. For amitraz, the adjusted LC_50_ value was 110 μg/g in 2003 [95% CI 39 – 217] and 12 μg/g [95% CI = 10-16] in 2022.

### Investigating mutations associated with pesticide resistance in mites

RNA-Seq data from a previous study comprised of Varroa samples from throughout New Zealand (Lester et al. 2022) was examined for gene mutations associated with pesticide resistance. None of the reads that mapped to the Varroa pyrethroid susceptible sodium channel gene (0-50 reads per sample, average 16.5) showed residue substitutions known to be associated with flumethrin resistance at amino acid positions 918 and 925. Similarly, in the Sanger sequencing analysis, the analysis of the 10 individual mite samples from across New Zealand showed no evidence of mutations in nucleotides 1710-1712 (amino acid position 925) (Supp. Table 1, Supplemental material). All mites presented the wild-type leucine residue at this position.

## Discussion

Colony loss rates over winter rose significantly in New Zealand between 2017 and 2021. During this same time period, attributions of losses to Varroa increased sharply, with beekeepers reporting this parasite to be the biggest driver of colony loss during winter 2021. These findings are unsurprising as Varroa has also been found to be the main cause of winter colony losses for countries such as the United States (Seitz *et al*. 2016, Kulhanek *et al*. 2017, Steinhauer *et al*. 2021). As honey exports are of great economic value in New Zealand, worth $482 million in 2021 (Stahlmann-Brown *et al*. 2022b), it would be beneficial for the apiculture industry to better understand why colony losses to Varroa have increased. Whilst the majority of commercial beekeepers were satisfied with the efficacy of flumethrin, approximately 19% of beekeepers in the survey indicated that flumethrin had failed to successfully control Varroa in their hives. Failure to control Varroa is consistent with emerging miticide resistance, although Stahlmann-Brown and Robertson (2022) report that significantly underdosing flumethrin was a common practice by New Zealand beekeepers during the 2020-2021 season. Even so, some commercial beekeepers communicated with the study authors that they felt flumethrin had become less effective in recent years; many of these beekeepers reported that they followed flumethrin treatments with other treatments such as oxalic acid. Indeed, treatment with oxalic acid has risen during the study period, and it is noted that oxalic acid can be applied during the honey flow if a spring treatment fails. However, oxalic acid may often not be ideal either as sub-lethal stress effects to honey bees have been reported (Gunes *et al*. 2017, Rademacher *et al*. 2017).

Our experiments testing for flumethrin resistance in New Zealand Varroa populations found evidence of resistance when compared to the study conducted by Goodwin *et al*. (2005). The adjusted LC_50_ for flumethrin (156μg/g) was found to be 13 times what it was in 2003 (12μg/g). However, issues with the variability of results from 2003 mean that a degree of caution is needed when drawing comparisons between the two studies. There was far greater variation in mite mortality in the 2003 study than was observed in 2022, particularly in the replicates for the lower concentrations of flumethrin. Preliminary data for the 2022 study found subtle differences in temperature and humidity affected mortality, which is a possible explanation for the variation observed in the 2003 study. In the 2022 trials, mortality rates were consistent throughout.

The methodology used by Goodwin *et al*. (2005) was based on a study conducted by Milani (1995) assessing the susceptibility of Varroa to flumethrin, fluvalinate and acrinathrin in Italy. This study also dissolved chemicals into paraffin wax, making their results comparable to the current study. For the flumethrin trials, the LC_50_ values observed for two non-resistant mite populations in the Italian study were 0.28 μg/g and 0.36 μg/g, substantially lower than the LC_50_ of 156 μg/g we observed in 2022. Additionally, the LC_50_ values in our current study were seven times greater than the LC_50_ values (11.4 μg/g and 20.5 μg/g) for the two flumethrin-resistant Varroa populations in Italy (Milani 1995). This difference further suggests that Varroa populations in New Zealand are likely to have developed a degree of resistance to flumethrin.

In contrast, trials assessing the efficacy of amitraz found no evidence of mites developing resistance since 2003. In fact, the estimated LC_50_ value for amitraz in the current study (12 μg/g) was much lower than in 2003 (110 μg/g, Goodwin *et al*. (2005)). The apparent drop in LC_50_ value is likely not due to mites becoming more susceptible to the treatment, but was likely due to issues with the 2003 trials which led to the unexpected survival of mites at higher concentrations of amitraz. In 2003 the average mite mortality rate did not exceed 65%, even for the highest concentration of 300 μg/g, which far exceeds the maximum concentration tested in other studies that were able to achieve 100% mortality (Thompson *et al*. 2002, Maggi *et al*. 2008). At the time, it was suggested that this unexpected result was due to the chemical not being sufficiently mixed into the wax. It may be possible that the mites in Goodwin *et al*. (2005) had inconsistent exposure to the pesticide, or that there were other methodological issues with their setup. Their study, however, represents the best available data we have for comparison. Whilst there is some question about the amitraz results in 2003, the LC_50_ in 2022 suggests that amitraz is at least as effective as it was, so we are less concerned about the efficacy of this pesticide than we are for flumethrin.

The experiment conducted by Milani (1995) did not test for amitraz resistance, and other published studies of resistance to this chemical used different methodologies such as direct exposure of the mites to treatment strips or vials of evaporated solution rather than amitraz-impregnated paraffin wax (Milani 1995, Thompson *et al*. 2002, Maggi *et al*. 2008). This prevents us from drawing comparisons to other findings but does provide a baseline for future studies. Examining the feedback from beekeepers that completed the New Zealand Colony Loss Survey, ~28% found amitraz to be completely successful in treating Varroa, and less than 10% found amitraz to be partly successful or not at all successful. Amitraz was thus considered more effective than flumethrin by commercial New Zealand beekeepers. It remains possible that undetected resistance to amitraz is developing in New Zealand mite populations as there has been evidence of resistance in other countries (Kamler *et al*. 2016, Almecija *et al*. 2020, Rinkevich 2020). However, for the most part, amitraz seems to still be a popular and effective treatment against Varroa in many countries, even after decades of use (Ferland *et al*. 2021, Hernández-Rodríguez *et al*. 2021).

Molecular analyses conducted on the pyrethroid-susceptible sodium channel gene in Varroa found no evidence of mutations known to be associated with flumethrin resistance. This result was surprising as numerous studies on Varroa populations with known resistance to pyrethroids have been observed to possess mutations within this gene (González-Cabrera *et al*. 2013, González-Cabrera *et al*. 2018). Evidence suggests that the resistance of Varroa to pyrethroids has only evolved once or twice, initially arising in mite populations from Italy before dispersing to other regions via the movement of bee colonies (Martin 2004, Mitton *et al*. 2022). Mitochondrial gene analysis of Varroa indicate only one introduction of this parasite into New Zealand (Lester *et al*. 2022). It is therefore likely that the Varroa introduced to New Zealand did not already possess known pyrethroid-resistant mutations. However, Varroa in New Zealand may exhibit novel mutations that would similarly confer flumethrin resistance.

Although further research is needed, the findings of our trials, in conjunction with reports from beekeepers, suggest mites may be developing resistance to one of the most popular Varroa treatments in New Zealand.

The high level of inbreeding involved in Varroa reproduction and haplo-diploid sex determination allows for the rapid fixation of beneficial mutations in a population (Beaurepaire *et al*. 2017, González-Cabrera *et al*. 2018). This ability of resistant genes to spread swiftly through a mite population is why it is so important to detect and attempt to mitigate miticide resistance early. The development of resistance to chemical treatments by Varroa highlights how crucial it is to develop new control strategies against Varroa. The detrimental effects these miticides have on the honey bees themselves are an additional motivator for new management approaches (Tihelka 2018). One new strategy currently being investigated is the breeding of Varroa resistant traits in honey bees, such as hygienic behaviour, grooming and shorter brood development times (Spivak and Gilliam 1998, van Alphen and Fernhout 2020). These approaches may be more sustainable than pesticides; however, there have been challenges in attempts to maintain mite resistant traits within bee populations due to the heritability of these traits, genetic variability within hives and a poor understanding of the combination of traits required to achieve natural resistance (Mondet *et al*. 2020). Another strategy currently under development which shows more promise is the utilisation of RNA interference technology (RNAi) against Varroa mites (Garbian *et al*. 2012). This method has been observed to reduce mite populations (Garbian *et al*. 2012, Huang *et al*. 2019) and is thought to be species-specific to Varroa, likely making it harmless to honey bees and other non-target species (Tan *et al*. 2016, Krishnan *et al*. 2021).

Current management strategies are providing a degree of protection for honey bee populations. However, there is a need for resistance management to ensure chemicals including flumethrin remain effective. Alternating mite control treatments helps prevent the development of resistance and is a management strategy that the majority of commercial beekeepers in New Zealand utilise according to the findings of our study. There is still concern that not all beekeepers are practicing correct resistance management, as 13% of beekeepers that participated in the 2021 survey (which included hobbyists) reported solely using flumethrin to treat for Varroa (Stahlmann-Brown *et al*. 2022a). The ability of mites to develop resistance to chemical treatments highlights the need for more effective Varroa control methods to protect honey bees, and to help prevent severe economic losses and threats to food security globally.

## Data Availability Statement

Data from the New Zealand Colony Loss Survey are not publicly available due to privacy concerns and potential commercial sensitivities. Data from the experiments testing for chemical resistance presented in this study is available on request from the corresponding author.

## Informed Consent Statement

The New Zealand Colony Loss Survey undergoes an annual social ethics review by Manaaki Whenua — Landcare Research following guidelines of the Code of Ethics developed by the New Zealand Association of Social Science Researchers.

