## Supplementary material for "Are increasing honey bee colony losses attributed to *Varroa destructor* in New Zealand driven by miticide resistance?": Figure S1, Supplemental material

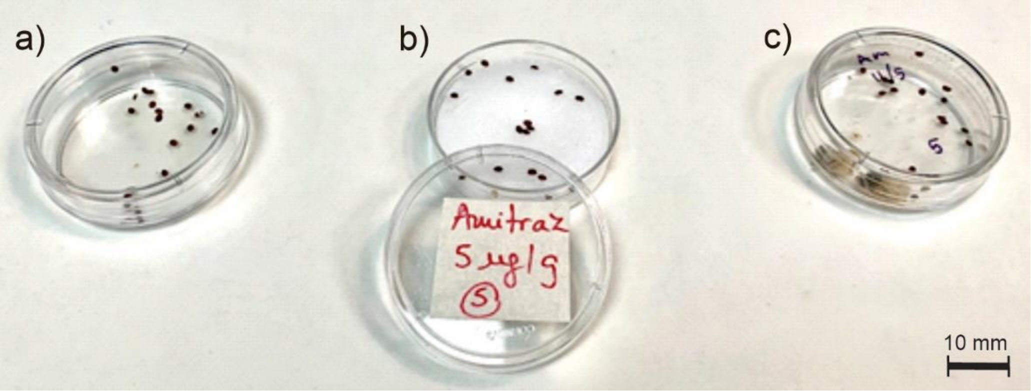


Figure S1. Experimental setup testing the effects of pesticide concentration on Varroa survival. a) 20 mites in a petri dish awaiting treatment, b) mites transferred to petri dish containing the treatment for an hour and c) mites in a new petri dish following exposure to the treatment with bee pupae for 48 hours, followed by a survival assessment. Photo credit: Phil Lester.

Supp. Table 1. Ten individual *Varroa destructor* mites from throughout New Zealand were sequenced in this study. We examined the samples for mutations in nucleotides 1710-1712 (amino acid position 925) of the coding region of the voltage-gated sodium channel (Na) gene (GenBank accession number KC152655) as determined in González-Cabrera et al. (2016). All our sequences presented the wild type allele.

| **Sample ID** | **Location** | **Sanger sequences using the Vd_L925V_F primer (nucleotides 1710-1712 are highlighted in red, and the Vd_L925_R primer binding region is highlighted in green).** |
|---|---|---|
| 1 | Hamilton (Matangi, Waikato) | ACGATAGGAGCTCTGGGTAACCTGACCTTTGTGTTGGGAATTATCATCTT |
| 2 | Tamahere (Waikato) | ACGATAGGAGCTCTGGGTAACCTGACCTTTGTGTTGGGAATTATCATCTT |
| 3 | Horotiu (Waikato) | ACGATAGGAGCTCTGGGTAACCTGACCTTTGTGTTGGGAATTATCATCTT |
| 4 | Enderly (Waikato) | ACGATAGGAGCTCTGGGTAACCTGACCTTTGTGTTGGGAATTATCATCTT |
| 5 | Ōtaki (Kapiti) | ACGATAGGAGCTCTGGGTAACCTGACCTTTGTGTTGGGAATTATCATCTT |
| 6 | Kelburn (Wellington) | ACGATAGGAGCTCTGGGTAACCTGACCTTTGTGTTGGGAATTATCATCTT |
| 7 | Vogeltown (Wellington) | ACGATAGGAGCTCTGGGTAACCTGACCTTTGTGTTGGGAATTATCATCTT |
| 8 | Ashburton (Canterbury) | ACGATAGGAGCTCTGGGTAACCTGACCTTTGTGTTGGGAATTATCATCTT |
| 9 | Ashburton (Canterbury) | ACGATAGGAGCTCTGGGTAACCTGACCTTTGTGTTGGGAATTATCATCTT |
| 10 | Ashburton (Canterbury) | ACGATAGGAGCTCTGGGTAACCTGACCTTTGTGTTGGGAATTATCATCTT |
